# Mindfulness meditation experience is associated with increased ability to monitor covert somatosensory attention

**DOI:** 10.1101/2020.02.04.933606

**Authors:** Stephen Whitmarsh, Ole Jensen, Henk Barendregt

## Abstract

The distinguishing practice of mindfulness meditation is the intentional regulation of attention towards the present moment. Mindfulness meditation therefore emphasizes metacognitive functions, in particular the ability to monitor the attentional focus on a moment-by-moment basis. In this study we set out to test whether mindfulness meditation experience is associated with an increased ability to monitor moment-by-moment fluctuation in the attentional state. In response to auditory cues, participants maintained somatosensory attention to either their left or right hand. At random moments, trials were terminated by a probe sound to which participants reported their level of attention at that moment. MEG was recorded during the attention interval preceding probe onset. Using a beamformer approach, alpha activity in contralateral primary somatosensory regions was quantified. Alpha activity for self-reported high versus low attention trials was compared both within and between groups of either highly experienced experienced mindfulness meditators, novice meditators or meditation-naive participants (controls). As predicted, generally contralateral alpha power was associated with self-reported attention. Novice meditators (< 1000 h of meditation) showed temporal profiles similar to controls, displaying a correspondence between self-report and alpha power preceding probe onset. Expert meditators (≫ 1000 h) showed a strikingly different pattern, however. Their self-reported attentional state corresponded with alpha power during a more extended time interval preceding those of controls and novice meditators. In addition, self-reported low attention trials showed a distinctive alpha suppression preceding probe onset, suggesting that the ability for moment-by-moment monitoring of the attentional state permitted greater attentional control.

## 1 Introduction

The goal of this study is to investigate whether extensive meditation experience can be associated with changes in meta-metacognitive monitoring of attention. While meditation practices are generally considered to incorporate the practice of attentional control, mindfulness meditation distinguishes itself by an emphasis on continuous metacognitive monitoring of mental contents and processes (Lutz, Slagter, Dunne, & Davidson, 2008; Lutz, Dunne, & Davidson, 2007). Mindfulness based interventions have been proven successful in treating a wide range of clinical syndromes ranging from depression (Chiesa & Serretti, 2011; Piet & Hougaard, 2011; van Aalderen et al., 2012) to anxiety (Roemer & Orsillo, 2006; Würtzen et al., 2013) and ADHD (Zylowska et al., 2008). Several clinical interventions have integrated mindfulness, e.g. in Mindfulness Based Stress Reduction (MBCT, (Kabat-Zinn, 2014, 2013), Mindfulness Based Cognitive Therapy ((Segal, Williams, & Teasdale, 2002; Teasdale, Segal, & Williams, 1995) and Acceptance and Commitment Therapy (Hayes, 2004). The mechanisms by which mindfulness meditation might benefit the mental health of the practitioner remain unclear, however. Initial proposals have been put forward from cognitive psychological (Baer, 2003; Brown, Ryan, & Creswell, 2007) or neuroscientific perspectives (e.g. (Chiesa & Serretti, 2011; Hölzel et al., 2011; Tang & Posner, 2013). in which mindfulness has been defined in terms of “paying attention in a particular way: on purpose, in the present moment, and nonjudgmentally” (Kabat-Zinn, 2014) or “the state of being attentive to and aware of what is taking place in the present” (Brown & Ryan, 2003) and similarly in (Bishop et al., 2004). The practice of mindfulness meditation is therefore understood beyond the ability to maintain attention upon a specific object or to safeguard oneself from (especially negative) distraction. Rather, mindfulness meditation emphases metacognitive awareness of experience regardless of its desirability or relevance. In fact, it is this explicit awareness of ongoing mental content, called meta-awareness (J. W. Schooler et al., 2011) or metacognition (Fleming, Dolan, & Frith, 2012), that allows distracting mental content to be noticed so that attention can subsequently be reoriented (J. W. Schooler et al., 2011; Smallwood, Beach, Schooler, & Handy, 2007; Smallwood & Schooler, 2006). Mindfulness, or “attending to the present moment”, can therefore be operationalized as the metacognitive monitoring of the focus of attention.

It is well known that mind-wandering often occurs without the awareness that one’s mind has drifted (Giambra, 1995; J. Schooler & Schreiber, 2004). Importantly, recent work has show that the tendency to mind-wander correlates negatively with a mindful disposition (Mrazek, Smallwood, & Schooler, 2012). Furthermore, successful outcome after MBCT has been associated with increased availability of metacognitive reflection on negative experiences (Teasdale et al., 2002). Although metacognition is understood to be an important aspect of mindfulness, experimental evidence for improved metacognitive functioning after mindfulness meditation has been only inferred indirectly from studies showing increased performance in tasks requiring cognitive control and response inhibition. Jha and colleagues (2007) found that mindfulness meditation improved conflict monitoring in the Attention Network task. Allen and colleagues (2012) showed a reduction in affective Stroop conflict and in Heeren and colleagues (2009), mindfulness training was associated with increased cognitive inhibition in go/nogo task. Metacognition has been a traditional topic of interest in memory research for decades (see: (Metcalfe & Shimamura, 1996) and (Dunlosky & Bjork, 2014) for an overview). Recently, metacognition of action and perceptual processes have become a topic of interest as well (see (Fleming et al., 2012) for an overview). Since mindfulness meditation is first and foremost an attentional practice, recent work on metacognition of attention, or conversely of mind-wandering, could be a promising direction of neuroscientific research. Previous experiments on metacognition of attention or mind-wandering have used control tasks or external stimulation (Braboszcz & Delorme, 2011; Christoff, Gordon, Smallwood, Smith, & Schooler, 2009; Macdonald, Mathan, & Yeung, 2011; Wilimzig, Tsuchiya, Fahle, Einhäuser, & Koch, 2008). When studying neural correlates of attention during meditation, however, the performance of a concomitant task can be expected to interfere with attentional monitoring, as well as preventing ecologically valid conclusions regarding meditation. Furthermore, task performance and stimulus processing might provide information about the attentional state during stimulus processing or task performance, preventing conclusions in terms of metacognition of attention *per se* (Fleming et al., 2012). Importantly however, the attentional state can be determined without a control task or external stimulus. Cortical alpha activity has been shown to reflect both the degree and the location of covert somatosensory attention (Haegens, Handel, & Jensen, 2011; Haegens, Luther, & Jensen, 2012; van Ede, de Lange, Jensen, & Maris, 2011; van Ede, Jensen, & Maris, 2010). Furthermore, occipital alpha power was shown to be strongly correlated with self-reported attention during a visual detection task (Macdonald et al., 2011). We have reported that contralateral alpha power corresponds to self-reported attentional focus as well (Whitmarsh, Barendregt, Schoffelen, & Jensen, 2014; Whitmarsh, Oostenveld, Almeida, & Lundqvist, 2017). In this study we used an identical paradigm as in Whitmarsh et al. (2014) to investigate whether metacognitive awareness of attention is improved after extensive mindfulness meditation. Participants were presented with auditory cues to attend either to their left or right hand while their brain activity was measured using MEG. After a random time interval trials were terminated with a probe sound to which they reported the degree of attentional focus. Self-reported attentional focus was compared with contralateral somatosensory alpha preceding probe onset, in eleven one second intervals. We compared the degree of correspondence over time between novice and expert mindfulness meditators as well as with that previously reported for meditation-naive participants (Whitmarsh et al., 2014). We predicted that meditators would show a stronger and more sustained correspondence between self-reported attentional focus and contralateral alpha power over time. Furthermore, we expected these differences to be more pronounced for expert meditators compared to novices.

## 2 Method

### 2.1 Participants

The control group consisted of fifteen healthy participants (9 female, mean = 29.4 years, SD = 10.4). One control participant was excluded from the analysis due to excessive movement artifacts. The meditation group consisted of sixteen healthy experienced mindfulness meditators (5 female, mean = 48.0 years, SD = 16.1). They were recruited via personal network of mindfulness teachers or contacted directly. To be included in our study they had to be familiar with mindfulness meditation through a meditation retreat and had to practice mindfulness mediation regularly for at least a year at the time of the study. Participants were enrolled after providing written informed consent and were paid in accordance with guidelines of the local ethics committee (CMO Committee on Research Involving Humans subjects, region Arnhem-Nijmegen, the Netherlands). The experiment was in compliance with national legislation as well as the code of ethical principles (Declaration of Helsinki).

### 2.2 Procedure

Participants were instructed to attend continuously to the cued hand while remaining aware of their attentive state until a probe sound (2000 Hz tone) was presented (Figure 1A). Cues consisted of two sequential tones of 400 ms each, 200 ms apart, either ascending in pitch for the right hand (2000 Hz followed by 2500 Hz) or descending for the left hand (2000 Hz followed by 1500 Hz). Cue side was determined pseudo-randomly. The subjective evaluation of attention was done by button press, selecting one out of four options: (1) not at all, (2) little, (3) much, (4) fully/maximally attentive. To minimize head movements and provide more comfort participants were measured in supine position. To minimize eye movements and blinks and increase the chance of fluctuations in attentional focus, participants were instructed to remain with their eyes closed throughout the experiment. The experiment started with a training session until participants were familiar with the paradigm, followed by three continuous experimental blocks of 125 trials (≈ 25 min each) separated by self-paced breaks. Non-evaluation trials (10% ad random) were indicated by the presentation of an extra third cue of similar pitch to the second. Participants were instructed that no evaluation of attention was needed at these trials, but to respond as quickly as possible with their index finger. On these trials only, participants received a short electrical stimulation of the cued hand after probe offset, to provide an additional functional localizer (which was not used in the analysis). Non-evaluation trials were discarded from all further analysis. Cue-probe intervals were generated according to an exponential distribution with mean of 3 s and a cut-off time of 27 s seconds, providing a flat hazard rate. In other words, the chance of the probe occurring after trial onset was held constant. A minimal cue-probe interval of 5 s was added, resulting in an average cue-probe interval of 8 s seconds, and maximal of 32 s.

**Figure 1:**
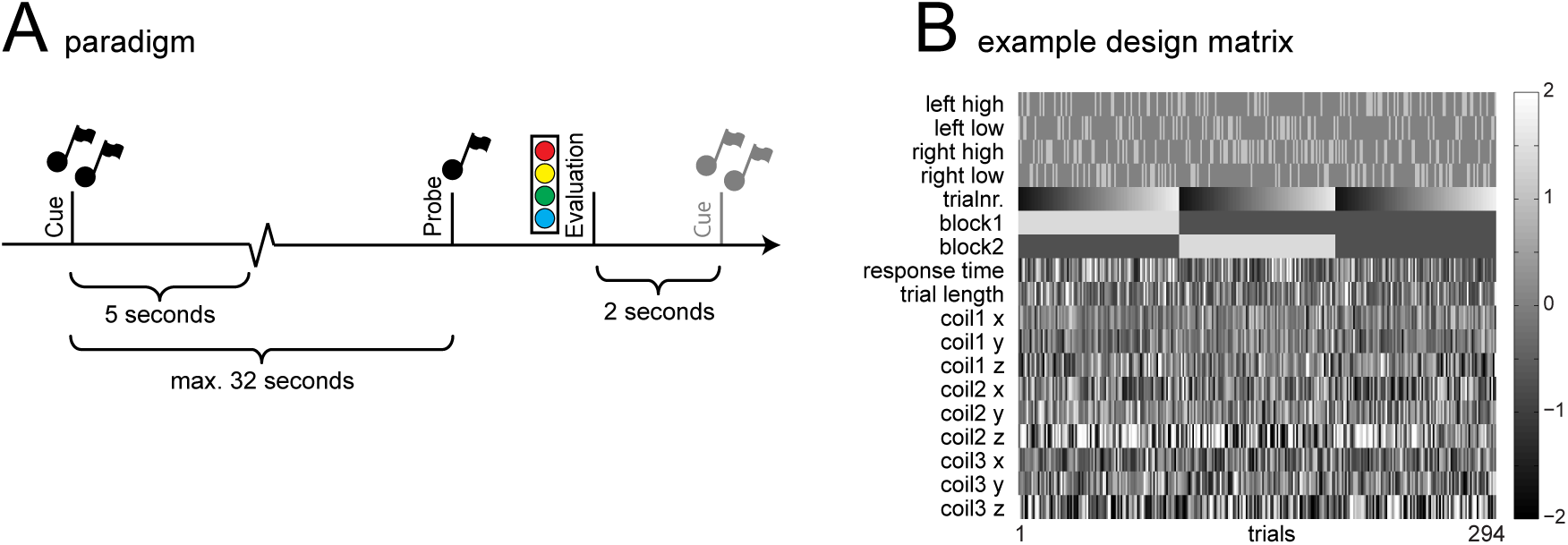
Schematic of paradigm and example of the design matrix used for the source level General Linear Model. A) Schematic depiction of paradigm showing timing parameters of trial. B) Example design matrix showing regressors for conditions (0’s and 1’s) and confound regressors (normalized).

### 2.3 Data Preprocessing

Continuous MEG data was recorded using a 275-sensor axial gradiometer system (CTF MEG TM Systems Inc., Port Coquitlam, BC, Canada) placed in a magnetically shielded room. The ongoing MEG signals were low-pass filtered at 300 Hz, digitized at 1200 Hz, and stored for off-line analysis. The subjects’ head position was continuously recorded relative to the gradiometer array using coils positioned at the subject’s nasion and at the left and right ear canals. High-resolution anatomical images (1 mm isotropic voxel size) were acquired using a 1.5 T Siemens Magnetom Sonata system (Erlangen, Germany). The same earplugs, using vitamin E instead of the coils, were used for co-registration with the MEG data.

### 2.4 Data Analysis

MEG data was analyzed using the Matlab-based Fieldtrip toolbox, developed at the Donders Institute for Brain, Cognition and Behavior (Oostenveld, Fries, Maris, & Schoffelen, 2011). Trials containing movement, muscle, and superconducting quantum interference device (SQUID) jumps were discarded by visual inspection. Independent component analysis (ICA) was used to remove eye and heart artifacts.

### 2.5 Source Reconstruction

Source reconstruction was performed using a frequency-domain beamformer approach (Dynamic Imaging of Coherent Sources) which uses adaptive spatial filters to localize power in the entire brain (Gross et al., 2001; Liljeström, Kujala, Jensen, & Salmelin, 2005). The brain volume of each individual subject was discretized to a grid with a 0.8 cm resolution. For every grid point a spatial filter was constructed from the cross-spectral density matrix and the lead field. The lead fields were calculated from a subject specific realistic single-shell model of the brain (Nolte, 2003), based on the individual anatomical MRIs. We calculated the cross-spectral density matrix based upon the full interval between cue offset and probe onset to obtain the most accurate estimation of the alpha sources. Individual alpha frequencies were used, determined by the maximum log power 7 Hz to 15 Hz on all trials and sensors.

Separate analyses were performed for one second segments preceding probe onset, to a maximum of 11 seconds before probe onset. A sufficient number of trials (≈ 100) had trial lengths of at least 11 seconds to enable source statistics at those intervals. After calculating the spatial filter for each grid point and one-second time segment, alpha activity was estimated using a (Slepian) multitaper approach to accomplish accurate frequency smoothing for the alpha band (±2 Hz) around the subject-specific alpha peaks. To enable valid voxel-by-voxel comparisons in the face of the beamformer depth bias, alpha estimates were standardized over trials.

A voxel-by-voxel first-level GLM approach was then used for every subject and time segment. Figure 1B shows an example design matrix, including cue side (left/right) and self-report (high/low), dichotomized according to a median split per subject and time-point. In addition, trial number as well as the mean X, Y and Z position of the three fiducial coils were entered as separate regressors. The locations of the fiducial coils indicate the position of anatomical landmarks of the subject’s head (nasion and pre-auricular points) in the MEG helmet. By including the average fiducial positions during each trial, variance that was caused by differences in head position over trials was also reduced. To further reduce variance that could be explained by response preparation, regressors for evaluation response times and cue-probe duration were added to the GLM, together with separate regressors for the response hand, which was switched between each block and randomized over subjects. The cue side and self-report predictors consisted of 0’s and 1’s, thereby yielding mean standardized alpha power after multiplication with the standardized data. By standardizing the remaining covariates (response time, trial length and fiducial position) multiplication with the standardized data resulted in correlation values (*ρ*). Prior to averaging and group statistics, the resulting beta-values and correlations values were spatially normalized using SPM2 to the International Consortium for Brain Mapping template (Montreal Neurological Institute, MNI, Montreal, QC, Canada).

### 2.6 Functional localization of somatosensory regions

After the reconstruction of alpha power for each voxel and time-point, somatosensory regions of interest (ROIs) were determined based on alpha power during the last second preceding probe onset. A voxel-by-voxel comparison was made between left and right attention trials. A cluster-based permutation test (Maris & Oostenveld, 2007) was then used to identify significant spatial clusters. This resulted in a distinct somatosensory alpha-ROI for each hemisphere. Each ROI therefore depended on cue condition (left versus right), but remained independent of the self-reported evaluation of attention.

### 2.7 Region of Interest analysis

Alpha power values within the left and right ROI voxels were averaged according to cue condition (ipsi versus contra), evaluation (high versus low) and time-point (eleven one-second intervals preceding probe onset). The effects of cue condition and evaluation on mean alpha power were tested over time using repeated measures ANOVA. Differences in these effects over time were tested using post-hoc t-tests per time-point.

## 3 Results

Attentional focus fluctuated over trials, as was reflected by the use of the full range of responses (Figure 2A). Participants were generally confident about their ability to perform the task, as was shown by the large fraction of high evaluations. A repeated measures ANOVA with one within-subject factor with four levels (evaluation) and one group factor (meditators vs. controls) showed that the number of trials per level of attention were significantly different (*F* (3) = 30.931, *p* < 0.001), and assumed a negative linear relationship (*F* (1) = 73.889, *p* =< 0.001). No effect of group on evaluation was found (*F* (3) = 1.451, *p* = .234). Evaluation times were also significantly different between levels of attention (*F* (3) = 55.390, *p* < 0.001) and showed a positive linear relationship (*F* (1) = 69.862, *p* < 0.001). No effect of group on evaluation time was found (*F* (3) = 2.038, *p* = .116). Furthermore, evaluation times correlated negatively with cue-probe duration for both controls 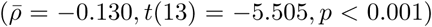 and meditators 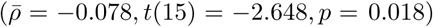, showing that longer cue-probe durations did not result in a loss of vigilance. The correlation between evaluation times and cue-probe duration did not differ between the groups (*t*(28) = −1.3450, *p* = 0.189). The behavioral analysis therefore showed no differences between meditators and controls, and both groups were confident in their performance of the task.

**Figure 2:**
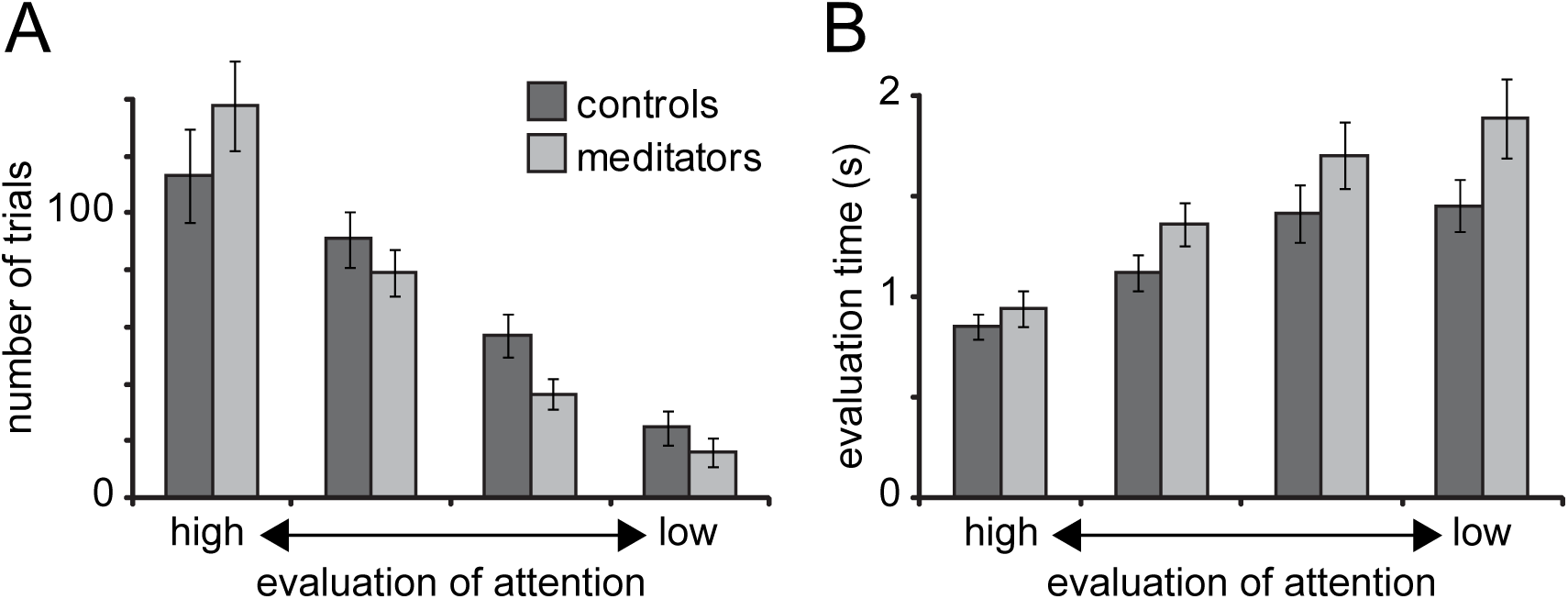
Behavioral differences between evaluations. A Distribution of number of trials per evaluation of attention show that attention was generally rated high, showing a negative relationship with level of attention. B Evaluation times were reduced when attention was evaluated higher, showing a positive relationship with level of attention. No differences between groups were found.

### 3.1 Regions of interest analysis

Somatosensory alpha regions of interest (ROIs) were determined for controls and meditators on the basis of the lateralization of alpha power (alpha cue left minus alpha cue right) during the last second before probe onset. A cluster-based permutation test (Maris & Oostenveld, 2007) was used to identify the significant clusters responsive to cue direction. In both groups the analysis resulted in two significant clusters, one in each hemisphere, including primary sensorimotor areas (see Figure 3). A cluster-based permutation test of alpha lateralization showed no significant differences between groups. Mean values of alpha power within the group-specific regions-of-interest were used in further analysis. We then computed contralateral alpha power preceding probe onset. Evaluations were dichotomized according to a median split per subject and time-point, resulting in time courses for high versus low attention trials (Figure 4).

**Figure 3:**
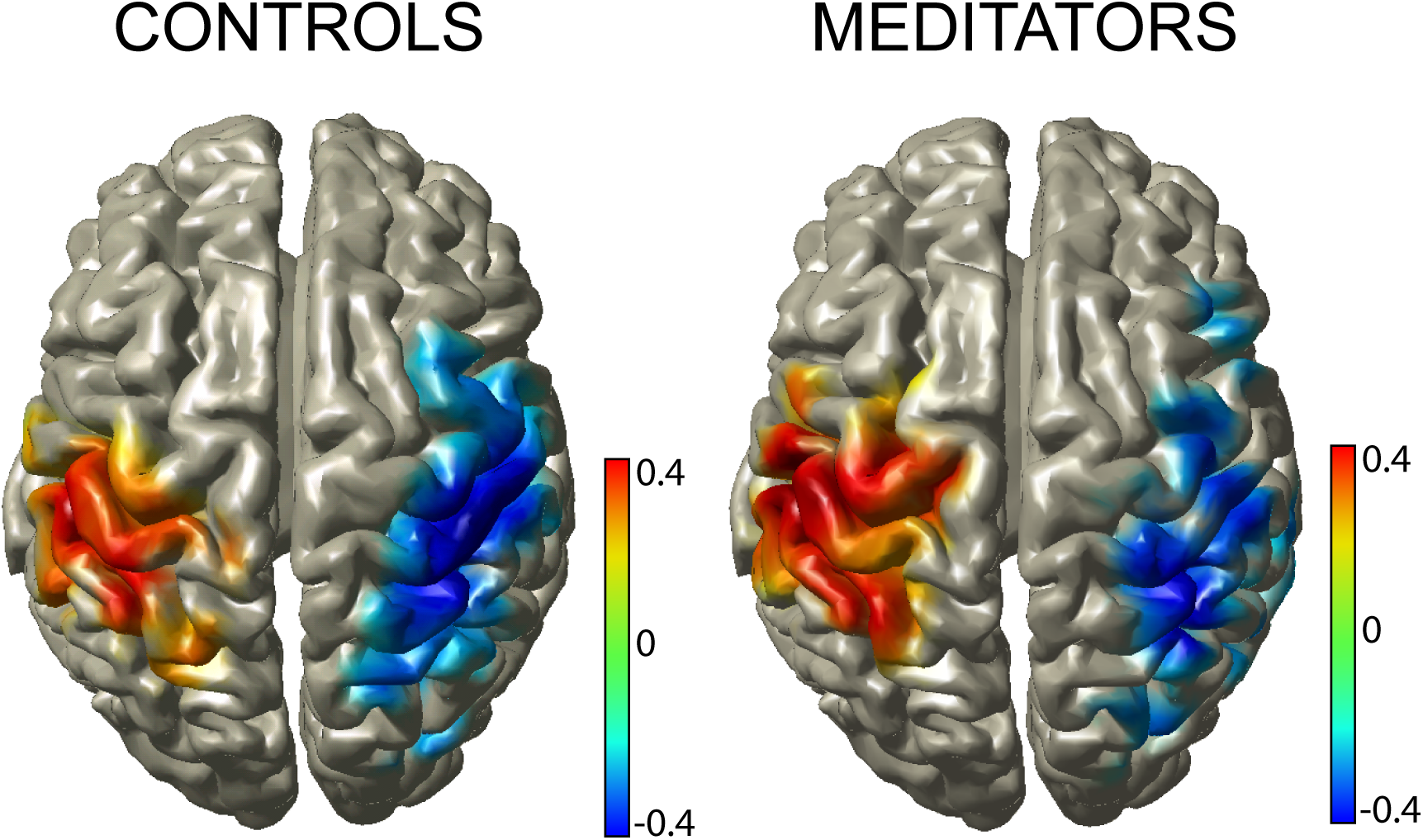
Somatosensory alpha power lateralized in response to cue direction in both groups. Source reconstructed alpha activity during one second preceding probe-onset shows clear lateralization in areas including primary somatosensory regions. Significant voxels were thresholded based on cluster-based permutation test (p¡ 0.05) (Maris & Oostenveld, 2007)

**Figure 4:**
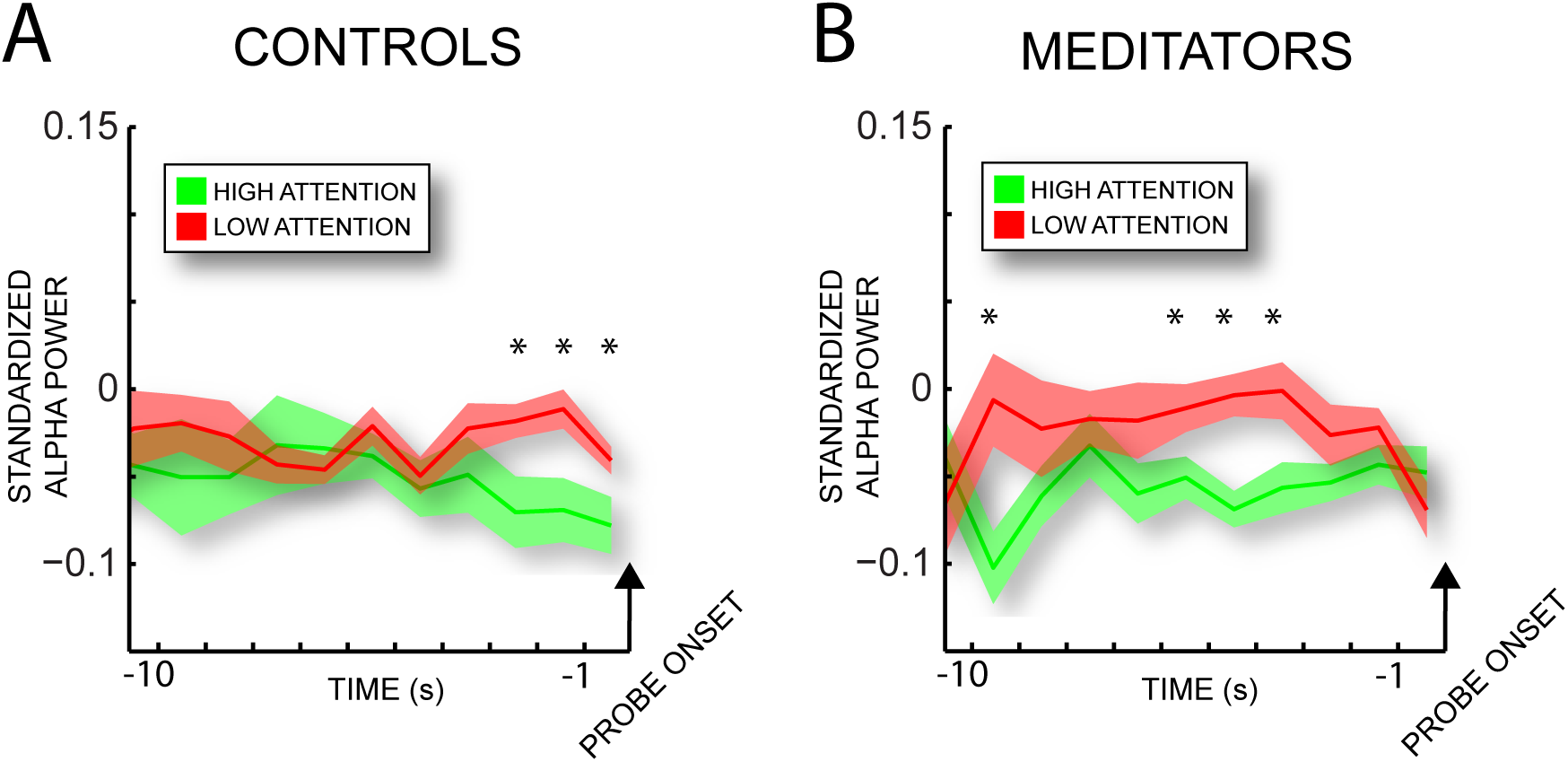
Time-resolved analysis of contralateral alpha power for controls (A) and meditators (B). Asterisk depict significant differences between high- and low-attention at individual time points. Shaded surface represents standard error of mean.

Differences between high and low attention trials were shown to be uncorrelated with age at any of the significant time points for either controls ([−3 : −2], *ρ* = −.31, *p* = .915; [−2 : −1], *ρ* = −.016, *p* = .956; [−1 : 0], *ρ* = .313, *p* = .275) or meditators: ([−10 : −9], *ρ* = −.912, *p* = .476; [−6 : −5], *ρ* = .096, *p* = .724; [−5 : −4], *ρ* = −.191, *p* = .478; [−4 : −3], *ρ* = .188, *p* = .486).

### 3.2 Controls versus meditators

We performed a repeated measures ANOVA with time and self-report (high vs. low) as within-subject factors and group (meditators vs. controls) as a between-subjects factor. Self-report showed a significant effect on contralateral alpha power (*F* (1, 28) = 9.509, *p* = 0.005). Time showed to be a significant factor as well (*F* (10, 19) = 2.983, *p* = 0.019). No significant interaction with group or between time and self-report effects were found. As expected, no effect of self-report was found for the ipsilateral ROI (*F* (1, 28) = 2.318, *p* = 0.139). We took a closer look at the temporal profile of alpha in the high versus low evaluated trials. In controls, the last three second interval before probe onset showed significantly lower alpha power in high versus low attention trials (Figure 4A; paired t-tests, two-tailed: [−3 : −2], *t*(13) = −2.357, *p* = 0.036; [−2 : −1], *t*(13) = −2.746, 0.017; [−1 : 0], *t*(13) = −2.648, *p* = 0.020). Although the full model ANOVA did not show a significant interaction between time and group, differences in alpha power preceded those of controls (paired t-tests, two-tailed: [−10 : −9], *t*(15) = −2.806, *p* = 0.04; [−6 : −5], *t*(15) = −2.156, *p* = 0.048; [−5 : −4]*t*(15) = −4.413, *p* < 0.001; [−4 : −3], *t*(15) = −2.275, *p* = 0.036). As predicted, no time points showed a difference between high and low attention on the ipsilateral ROI, for either the control group or the meditation group.

### 3.3 Novice versus expert meditators

To explore differences between novices and expert meditators we applied a median split of the meditation group on the basis of total hours of meditation experience. Hours of meditation experience were calculated from the self-reported estimated time spend in meditation per week, multiplied by the total duration of consistent meditation practice. For meditation retreats an estimated ten hours per day were added. The median split resulted in two very separate groups for novices (n = 7, mean = 555 hrs, range 220-885 hrs) and expert meditators (n = 8, mean = 12962 hrs, range 3275-24819 hrs). One meditator did not reply to our follow-up questions and could not be included in the analysis. As can be seen in Figure 5A, the temporal profile of the novice meditators showed a remarkable similarity with those of controls, showing a significant effect of self-report at a distinct period directly preceding probe onset ([−2 : −1]*t*(6) = −4.057, *p* = .006; [−5 : −4]*t*(6) = −2.298, *p* = .062; [−6 : −5]*t*(6) = −3.865, *p* = 0.008). Expert meditators, however, showed a dramatically different profile (Figure 5B), with significant differences in alpha power preceding the differences found in novices ([−10 : −9]*t*(7) = −2.391, *p* = 0.048; [−8 : −7]*t*(7) = −2.201, *p* = 0.064; [−7 : −6]*t*(7) = −2.008, *p* = 0.085; [−5 : −6]*t*(7) = −2.580, *p* = 0.036; [−5 : −4]*t*(7) = −3.769, *p* = 0.008). Furthermore, the last interval in high meditators showed a significant difference in the opposite direction after what seems to be a gradual reduction of alpha power in low attention trials during the last seconds of the trial (*t*(7) = 3.593, *p* = 0.009). We therefore divided the temporal profile midway in time and averaged alpha power for high and low self-report within the late and early period preceding probe onset. An ANOVA with self-report (high vs. low) and time (early vs. late) as within-subjects factors and group (novice vs. expert) as between-subjects factor showed a significant main effect of self-report (*F* (1, 13) = 7.672, *p* = 0.016) as well as significant interaction between self-report, time and group (*F* (1, 13) = 8.908, *p* = 0.011).

**Figure 5:**
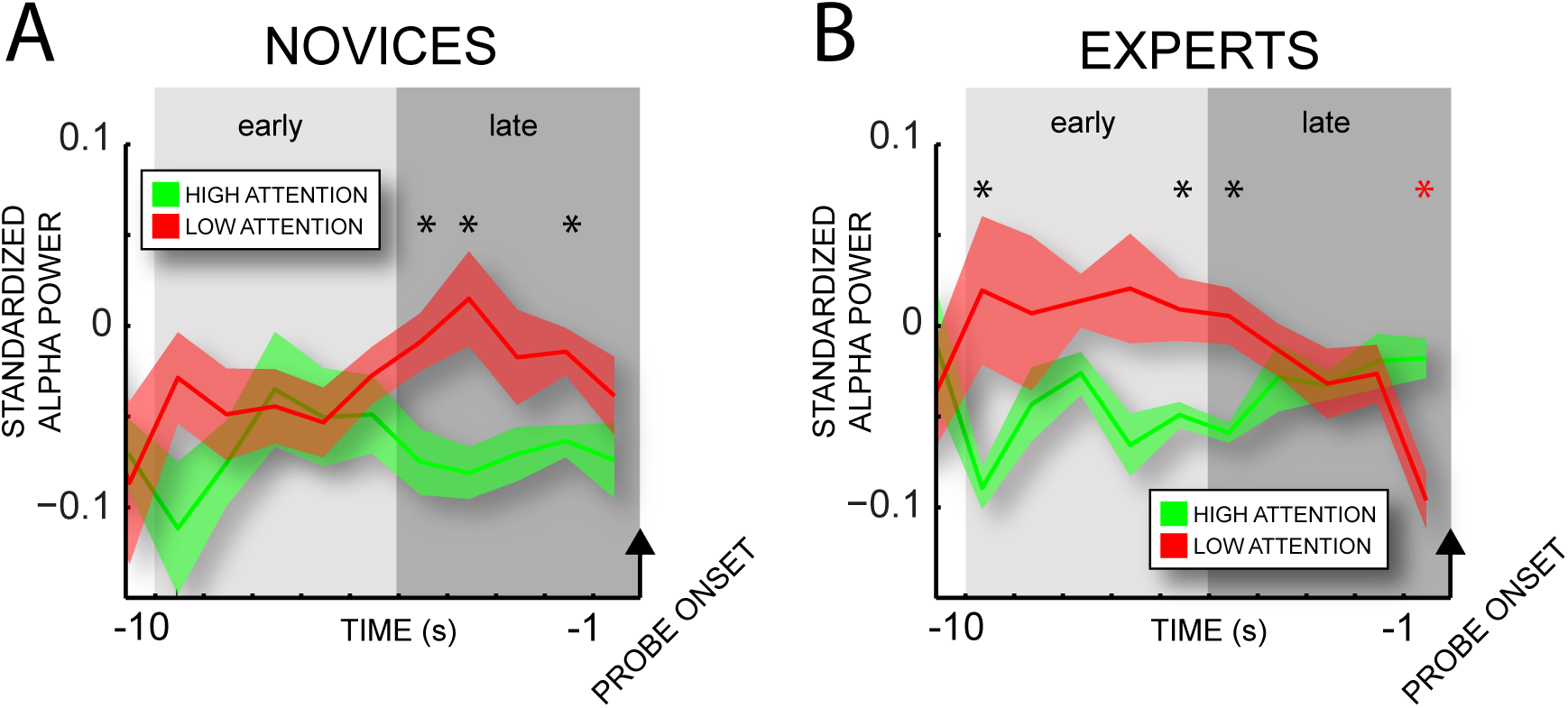
Time-resolved analysis of contralateral alpha power for novice meditators (A) and expert meditators (B). Asterisk depict significant differences between high- and low-attention at individual time points. Shaded surface represents standard error of mean.

## 4 Discussion

Compared to both meditation-naive controls and novice meditators, expert mindfulness meditators showed a distinctly different temporal profile of metacognitive awareness. While the self-reported attentional state of controls and novices were reflected by differences in alpha power preceding probe-onset, expert meditators showed a correspondence between self-report and alpha power during an extended period of time preceding that of both controls and novice meditators. Furthermore, expert meditators displayed a pronounced suppression of contralateral alpha power before probe onset, even surpassing those of high attention trials. Both these results are unexpected and warrant only tentative conclusions. Given these reservations, the results suggest shed light on previously unexplored territory and suggest new avenues of exploring neurocognitive mechanisms of mindfulness meditation. Firstly, the findings suggest that while novice meditators and controls were able to report on their attentional state retrospectively (i.e. after probe onset), expert meditators monitored their attention continuously and used it proactively rather than reactively. The ability maintain moment-by-moment evaluation of attention is in accordance with the proposed aim of mindfulness meditation. This might have enabled expert meditators to reestablish their attentional focus when distraction was noticed, as reflected by the suppression of alpha power in the latter half of low attention trials. However, since these trials were still reported as low attention trials, expert meditators might have confounded metacognitive awareness of their attentional state with that of attentional control. Future research could test this hypothesis by using additional self-report probes of both attentional state as well as effort/control. Positive findings on both indexes would provide objective evidence for the claim that mindfulness meditation increases the ability to differentiate internal cognitive processes (Bishop et al., 2004).

Alpha values for each voxel and time interval were standardized over all trials to remove the beamformer depth bias. Because we were interested in the correspondence between self-reports and alpha power over time, our data was standardized per time-interval as well. This made our paradigm particularly sensitive to metacognitive awareness of attention, but might have missed other effect of mindfulness meditation such as an increased ability to modulate somatosensory alpha (Kerr et al., 2011). Our findings are, however, in line with recent studies reporting improvements in attentional stability (Lutz et al., 2009) and improved conflict monitoring (Jha et al., 2007) as both are considered to be under metacognitive control (Fleming et al., 2012). The experiment required a minimum of behavioral responses, and these were not experienced as interfering with the usual practice of mindfulness meditation. In fact, the paradigm was well received by the expert meditators we measured. The benefit of a less intrusive paradigm in meditation research cannot be overstated. The absence of any external stimuli or behavioral measurements too often precludes unequivocal interpretation of psychophysical findings (see e.g. (Cahn & Polich, 2006)), while at the same time, exogenous perceptual and behavioral tasks might interfere with the introspective practice.

A possible limitation to our study is the fact that the meditation and control groups were not matched in age, with the meditation group being significantly older. However, for both controls and meditators, age did not correlate with the differences in alpha power between high and low attention trials. It is therefore highly unlikely that results were dependent on age. As a between-subjects cross-sectional study our results might also suffer from a selection bias. The fact that our findings depended on the hours of meditation experience speaks against an interpretation in terms of differences preceding their meditation practice. Lastly, we cannot exclude any ‘third variable’ that might correlate both with meditation experience as well as metacognitive ability. For instance, it is shown that metacognition is impaired under conditions such as cigarette craving (Sayette, Schooler, & Reichle, 2010) or inebriation (Sayette, Reichle, & Schooler, 2009). Since meditation is often practiced in a social and religious context, it cannot be excluded that influences of life-style might have played a role.

Mindfulness meditation has been theorized to distinguish itself from meditation practices that focus on sustained and selective attention through its emphasis on metacognitive monitoring (Lutz, Dunne, & Davidson, 2006; Lutz et al., 2008). While there is growing evidence that mindfulness meditation improves attentional abilities, convincing evidence for improved ability for metacognitive monitoring has been lacking (Chiesa & Serretti, 2011). This is unfortunate, since these metacognitive abilities are proposed to underlie its clinical efficacy (Baer, 2003; Teasdale et al., 2002). Our findings therefore provide important preliminary evidence for the involvement of metacognitive monitoring in mindfulness meditation. Furthermore, our results suggest that such proactive attentional monitoring abilities could improve the ability to manage attentional resources on a moment-by-moment basis. However, in line with the conclusion of Chiesa and colleagues (2011), our results also show that measurable effects might require extensive practice.

## 5 Acknowledgements

We would like to express our gratitude to David A. Holmes for his invaluable insights in understanding the practices and potential of mindfulness meditation, and for his help in proofreading the manuscript. The authors also gratefully acknowledge the support of the BrainGain Smart Mix Programme of the Netherlands Ministry of Economic Affairs and the Netherlands Ministry of Education, Culture and Science the Netherlands Initiative Brain and Cognition, a part of the Organization for Scientific Research (NWO) under grant number SSM0611,

